# BatSpot: a retrainable neural network for automatic detection and classification of bat echolocation and detection of buzzes and social calls

**DOI:** 10.64898/2026.03.11.711063

**Authors:** Simeon Q. Smeele, Christopher Hauer, Christian Bergler, Dina K. N. Dechmann, Melina T. Dietzer, Morten Elmeros, Esben T. Fjederholt, Andrea Fogato, Jenna E. Kohles, Elmar Nöth, Signe M. M. Brinkløv

## Abstract

1. Bats are a diverse taxonomic group that display a wide range of interesting behaviours. Many bats are keystone species for their ecosystem, are IUCN Red-listed as vulnerable to critically endangered, and subject to human-wildlife conflicts arising from anthropogenic expansion. Yet bats remain understudied both with respect to behaviour, population ecology and conservation status. One of the major challenges when studying bats is obtaining data. Their nocturnal lifestyle and use of ultrasonic echolocation makes them difficult to track and record using traditional methods. Recent advances in passive acoustic monitoring have allowed researchers to record large amounts of data, but the detection and classification of vocalisations remain a challenge. Most available tools are either for profit or are limited to a narrow geographic range, and mostly focus on echolocation search phase calls.
2. Here we present BatSpot, a convolutional neural network trained to detect search phase calls, buzzes and social calls. It also offers the option to classify the search phase calls to species(-complex) level. We provide a GUI that allows researchers to retrain or transfer-train the models for their specific needs and validate the performance.
3. We test the performance of all models and show that they perform better than both commercial and open-source solutions (search phase file level F1: 0.97 vs 0.96, buzz detector F1: 0.95 vs 0.11). We furthermore show that retraining the search phase call detector for a new country with examples from just 59 recordings massively improves the performance (F1: 0.48 to 0.79).
4. BatSpot will enable bat researchers globally to automate detection and classification with minimal effort and includes novel options for social call and buzz detection, typically not featured in other automated tools for bat monitoring.

## Introduction

Bats are the second most diverse group of mammals (Burgin et al., 2018) with 1,500 species described (Simmons & Cirranello, 2020), and display a wide variety of complex vocal behaviours. They actively sense their surroundings using echolocation during both individual and social foraging (Jones & Holderied, 2007; Petrites et al., 2009; Egert-Berg & Hurme et al., 2018; Krivoruchkoetal et al., 2024), and display complex vocalizations related to social structure and mating (Knörnschild et al., 2017; Knörnschild et al., 2019; Smotherman et al., 2016). Bats are keystone species that provide important ecological services such as pest control, soil fertilisation, seed dispersal and flower pollination (Ghanem & Voigt, 2012; Ancillotto et al., 2017;Ramírez-fráncel et al., 2021). Loss of bat ecosystem services can have direct and profound effects on human health (Frank, 2024). Globally, numerous bat species are currently threatened by extinction (IUCN, 2024) and all European bat species are listed on Annex IV of the European Union’s Habitats Directive (92/43/EEC), requiring member states to monitor them and ensure favourable population status. There is increasing evidence that migrating bat species face increased mortality from expanding wind farms (Arnett et al., 2015; Voigt et al., 2022), as countries around the world have to balance the conservation of flying species against the need to shift energy production to renewable sources. Other threats that bat species face include climate change (Sherwin et al., 2012), disease (Hoyt et al., 2021) and declining insect populations (Forister et al., 2019). Addressing these challenges requires the development of robust methods for monitoring bat behaviour, activity, and populations.

Recording behaviour of bats which fly in the dark is challenging and monitoring population sizes becomes especially difficult for bats that roost and breed in cavities and potentially migrate long distances (Russo et al., 2018; Martínez-Fonseca et al., 2024; Rebolo et al., 2024). The most cost-efficient way to record bat activity on a large scale is the use of passive acoustic monitoring (PAM), which has become a well established method (Biffi et al., 2024, Roemer et al., 2025, Toshkova et al., 2025). Most bat species produce ultrasonic echolocation calls and social vocalisations that may span the audible and ultrasonic range. Echolocation calls can be assigned to three main categories. Search phase calls are usually species-specific but adaptable to the environment, with longer calls characteristic for open space foraging and commuting and shorter call duration typical in cluttered environments (Kalko and Schnitzler, 2001; Brinkløv et al., 2010). Approach calls are similar to calls produced in clutter, but might shift in frequency. The call interval (also called pulse interval) is also much reduced to yield quicker updates about prey or obstacle position. Finally, buzzes are produced during the very end of a prey interception, with much reduced call amplitude and call intervals. Social vocalisations are much more variable in some species and consist of longer multi-node sequences resembling bird song (Smotherman et al., 2016; Springall et al., 2019). Species can in some cases be recognised by the frequency range and modulation, the duration and the intervals between calls, while specific behaviours can sometimes be inferred from the type of call: buzzes mostly indicating feeding, social calls indicating varying social behaviours such as mate attraction, territorial defence and individual recognition (Knörnschild et al., 2017; Knörnschild et al., 2019; Smotherman et al., 2006).

While PAM is already effectively supplying vast amounts of audio data, the processing workflow often lags in efficiency. Approaches range from manual detection and classification of bat calls (Sugai et al., 2018) to more or less automated pipelines with or without manual validation of software performance. The applicability of commercial software is challenged by price and decreased transparency. Open source software also exists, but often comes trained for a limited geographic range focused outside the Global South, where bat species diversity is highest, and might not be easy to adapt and use without a computer science background. An exception to this is BatDetect2 (Aodha et al., 2022b), which contains pre-trained models for several countries, including two in the Global South. It does, however, require some familiarity running scripts from the console, especially if fine-tuning of the model is needed. For all these software, retrainability remains an issue. If used in a soundscape with new noise sources or bat species, performance is likely to drop. Retraining without coding experience is often not possible. Finally, all but one of the open source solutions have focused on search phase calls, with the exception of Jameson (2024), who developed a buzz detector: *buzzfindr* for Canadian species. This lack of tools to detect social calls and buzzes globally hinders bat research, since foraging or social activity are often important behavioural components in a monitoring context (Dietzer et al., 2024).

To address this, we have trained several versions of BatSpot, which is based on ANIMAL-SPOT, a convolutional neural network designed to detect and classify animal vocalisations (Bergler et al., 2022). We present detection models for bat echolocation search phase calls, buzzes and social calls, as well as a classification model for the search phase calls. We compare the performance of these models to several commercial and open source solutions on a validation dataset. We provide examples on how to validate the models for a new location and how to retrain the model with a reduced amount of training data. The model also allows for transfer learning, adding additional classes to the classification model. Finally, we provide a GUI (see Figure 1) and a README on how to train, retrain, transfer-train and validate models.

**Figure 1.**
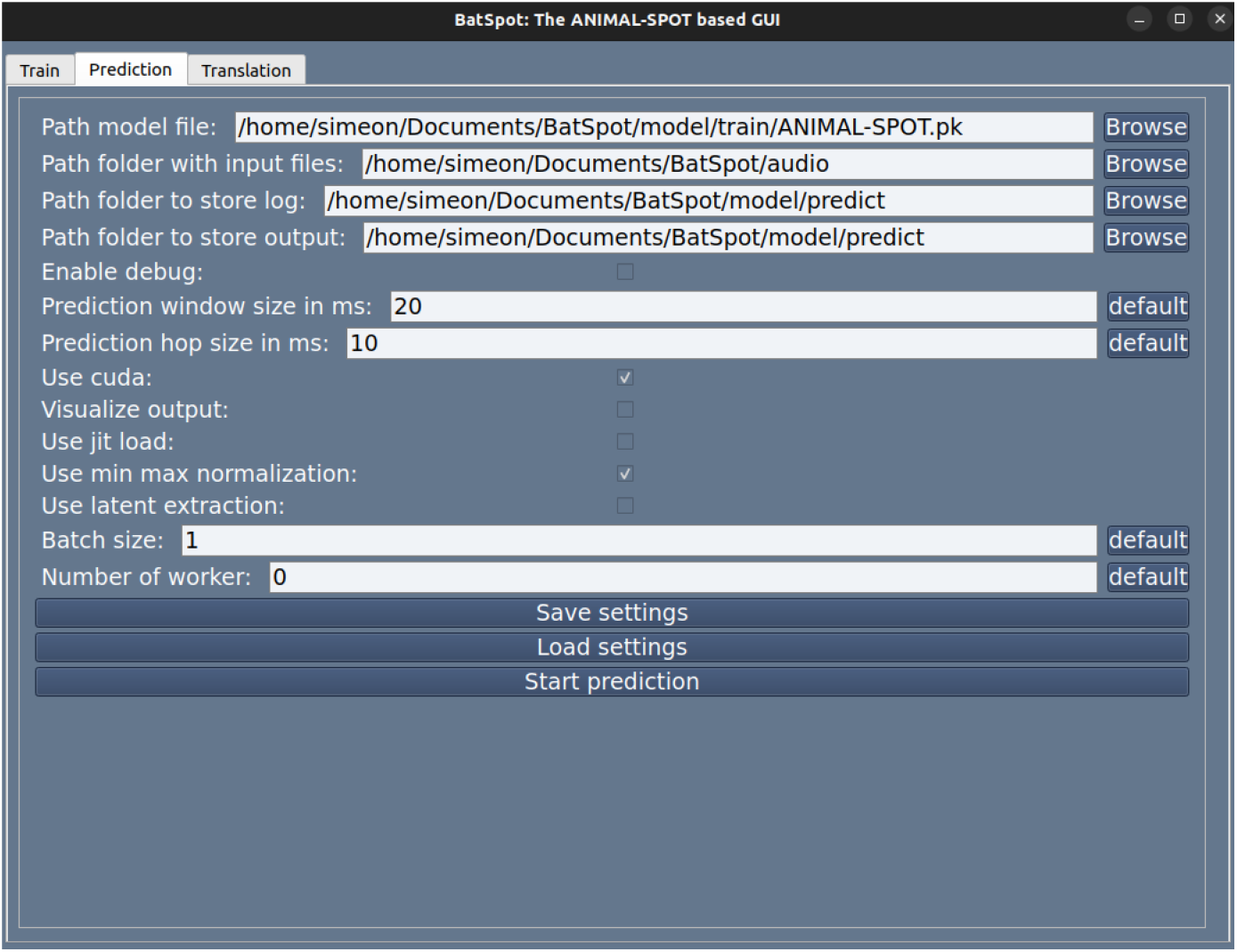
Screenshot of the GUI.

## Methods

### Call detector

To detect search phase calls in continuous recordings, we trained a binary version of BatSpot (hereafter called ‘call detector’). We used 12,396 noise examples and 10,766 target examples (search phase calls) from Denmark. We also used additional 2,000 noise examples for noise augmentation (these are added randomly in the background to increase generalisation ability). These were chosen randomly from the original noise set (with 14,396 noise examples). The model was trained on clips of 20 ms duration. Longer examples were randomly cut, shorter examples were zero-buffered. The frequency range was 1-95 kHz, this covers the high-energy part of all calls and is enough for detection. We report the validation and test accuracy, which is the proportion of true positives and true negatives out of the total number of validation/test examples. The reported validation accuracy is the highest accuracy during the training process, where the validation set is used to test if the model should stop training. The test accuracy is the accuracy on the test set, which is not shown to the model until after training has stopped.

To validate the model performance on full recordings we used 210 recordings from Denmark of variable duration. None of these had been used to generate training data. All search phase calls (including faint calls) were manually annotated. The call detector was run on windows of 20 ms, with 10 ms overlap between windows. The threshold for considering a detection true was 0.5. For each detection BatSpot gives a probability that it is correct in the prediction and by choosing 0.5 we allow only predictions where this probability is at least 0.5. Since the call detector and classifier are often used together, validation results are also presented together. For more detailed information, see below.

### Call classifier

To classify the detections from the call detector, we trained a multi-class version of BatSpot (hereafter called ‘call classifier’) including call examples of 12 species(-complexes). We used 1,161 examples of *Barbastella barbastellus*, 2,840 examples of *Eptesicus serotinus* (now known as *Chenaphaeus serotinus*), 1,217 examples of *Myotis brandtii/mystacinus*, 1,503 examples of *Myotis dasycneme*, 2,401 examples of *Myotis daubentonii*, 1,325 examples of *Myotis nattereri*, 2,397 examples of *Nyctalus noctula*, 982 examples of *Plecotus auritus*, 1,226 examples of *Pipistrellus nathusii*, 1,145 examples of *Pipistrellus pipistrellus*, 2,374 examples of *Pipistrellus pygmaeus* and 1,517 examples of *Vespertilio murinus*. Examples were drawn from the recordings made during the Danish national monitoring programme, where the focal individual is either observed in the field during recording or, if data quality allows, annotated by at least two experts. We also included 551 examples of buzzes, 899 examples of approach phase calls and 439 examples of social calls. These categories were not annotated to species level and considered as noise, since the call detector model was only supposed to detect search phase calls. To get optimal performance the approach phase call category was encoded as the noise category, serving as the default, if none of the other categories is predicted with high enough probability. The model was trained on clips of 20 ms duration. Longer examples were randomly cut, shorter examples were zero-buffered. The frequency range was 10-125 kHz, which meant that we had to use a slightly modified version of BatSpot, where the files were not downsampled to 192 kHz during processing, but rather to 250 kHz. This was done, since some species have calls extending beyond 96 kHz, information that can be used by the model during classification. We report the validation and test accuracy, which is the proportion of correct classifications out of the total number of validation/test examples.

To validate the model on detections from full recordings, we used the same validation set as for the call detector, and ran the call classifier on all the detections from the call detector. Since several *Myotis* species are notoriously difficult to discriminate from each other, we pooled both the validation and classifications from the model into the genus level label M. Depending on context, the same may be true for the low frequency species *E. serotinus, N. noctula* and *V. murinus*, thus these were pooled into an ENV complex. Calls assigned to *B. barbastellus* were considered noise, since they should not occur in the area where the validation data was collected. If calls could not be assigned to a species (-complex) or multiple calls were overlapping, these were excluded from validation. The call classifier was run on windows of 20 ms, with 15 ms overlap between windows. The overlap was increased to increase the chance of the classifier getting a window with the whole call included. The threshold for considering a detection true was 0.5. To quantify performance we created a confusion matrix and computed performance statistics. For the detection step we computed: (1) recall = tp/(tp+fn), which gives the fraction of calls that were detected out of the total number of calls; (2) precision = tp/(tp+fp), which gives the fraction of correct detections out of all detections; and (3) F1 = 2 * (recall*precision)/(recall+precision), which gives the average across the two; where tp = number of true positives, tn = number of true negatives, fp = number of false positives and fn = number of false negatives. For the classification step we computed: accuracy = correct classifications /(correct classifications + mistakes), which also included classifying the false positives of the detection model (where the correct classification was noise). For the overall performance, we computed: accuracy = correct classifications /(correct classifications + mistakes + misses), where the misses were the false negatives from the detection model. Since some projects only need to know if a species is present in a file or not, we also summarised the output per file by taking the species with most calls predicted and created a confusion matrix at file level, where we excluded 10 out of the 210 files, as they contained calls that could not be assigned within *Pipistrellus* (for the call level performance only these few calls were excluded and all other calls in the file were still used). We opted not to include area-under-the-curve values for either analysis, as these are harder to interpret.

To compare the combined performance of the BatSpot call detector and classifier to existing software, we ran the open source software BSG-BAT (Meramo et al., 2025), BatDetect2 (Aodha et al., 2022) and BAT (Fundel et al., 2023), as well as the commercial softwares Sonochiro (version 4.1.4, Biotope, France, biotope.fr), Kaleidoscope (version 5.8.1, Wildlife Acoustics Inc, USA, wildlifeacoustics.com) and BTO bioacoustics pipeline (*BTO Acoustic Pipeline*, n.d.). We used the same validation set as for the validation of BatSpot. In files with one or multiple calls that could not be assigned to a species (-complex), it would not be possible to assign a label at the file level. For each software we used default settings and followed best practise described in the manual. For BatDetect2, we changed the threshold for accepting a detection to 0.4 to balance the recall and precision. For each software, we compared the primary detection and classification to the ground truth (manual annotation by an expert) at file level. Where the software made multiple predictions, only the prediction with highest probability was used, since each of the validation files only contained a single species. If the software predicted species not present in the study area or did not classify the detection, the file was considered to contain only noise. For each software we show the confusion matrix and compute the recall, precision and F1 score.

### Buzz detector

To detect buzzes in continuous recordings, we trained a binary version of BatSpot. We used 2,293 noise examples and 1,288 target examples from Denmark, Germany and Panama. We also used 200 additional noise examples for noise augmentation. These were chosen randomly from the original noise set (of 1,488 noise examples). The model was trained on clips of 200 ms duration. Longer examples were randomly cut, shorter examples were zero-buffered. The frequency range was 1-95 kHz, which contains the part of the buzz with the most energy for most species. We report the validation and test accuracy, which is the proportion of true positives and true negatives out of the total number of validation/test examples.

To validate model performance we used 30 recordings from Denmark of variable duration (3-15 s), 100 recordings of 55 s from Germany and 90 recordings from Panama of variable duration (0.2-0.9 s). None of these had been used to generate training data. All buzzes (including faint and incomplete buzzes) were manually annotated. The buzz detector was run on windows of 200 ms, with 100 ms overlap between windows. The threshold for considering a detection true was 0.5 for Denmark and Panama, but was set to 0.9 for Germany to balance recall and precision. For validation we computed: recall = tp/(tp+fn), precision = tp/(tp+fp), F1 = 2 * (recall*precision)/(recall+precision), where tp = number of true positives, tn = number of true negatives, fp = number of false positives and fn = number of false negatives.

To compare our buzz detector to existing tools we ran the function *buzzfindr* from the R package *buzzfindr* (Jameson, 2024) on the same validation recordings using the default settings. We show the confusion matrix and compute the recall, precision and F1 score.

### Social call detector

To detect social calls in continuous recordings, we trained a binary version of BatSpot. We used 1,296 noise examples and 1,412 target examples from Denmark. We did not use noise augmentation. The model was trained on clips of 75 ms duration. Longer examples were randomly cut, shorter examples were zero-buffered. The frequency range was 1-60 kHz. We report the validation and test accuracy, which is the proportion of true positives and true negatives out of the total number of validation/test examples.

To validate the model performance we used 19 recordings from Denmark of variable duration. None of these had been used to generate training data. All social calls (including faint calls) were manually annotated. If a social call was not clearly a single unit (parts separated by ca. 50 ms), each part was separately annotated. The social call detector was run on windows of 75 ms, with 50 ms overlap between windows. The threshold for considering a detection true was 0.9. For validation we computed: recall = tp/(tp+fn), precision = tp/(tp+fp), F1 = 2 * (recall*precision)/(recall+precision), where tp = number of true positives, tn = number of true negatives, fp = number of false positives and fn = number of false negatives.

### Retraining

Retraining is the use of a previously trained model to speed up and/or improve performance of a new model. For retraining, the weights of the old model are imported after which new training examples are used to (slightly) update these weights. To illustrate how retraining increases the performance, we ran the existing call detector on a validation set of 24 files of 5 s each from Germany. We then created a training set (using different recordings) consisting of 290 noise examples and 477 target examples from Germany. We also used an additional 50 noise examples for noise augmentation. These were chosen randomly from the original noise set (of 340 noise examples). With that we first trained a model from scratch using the same settings and then trained a model using retraining from the Danish call detector model. We restricted the number of learning rounds (epochs) to 100, but did not change any other settings. To compare we present the recall, precision and F1 for all three models (baseline, retrained and trained from scratch).

## Results

### Call detector and classifier

After 150 epochs (the maximum) the call detector had a validation accuracy of 0.98 and a test accuracy of 0.97. The call classifier trained for 81 epochs (after no improvement for 20 epochs) and achieved a validation accuracy of 0.88 and a test accuracy of 0.87. For validation of the models on full recordings, we annotated 6,477 search phase calls. Of these, 40 could not be assigned a species, 74 were overlapping, 193 were either *P. nathusii* or *P. pipistrellus* and 490 were either *P. pipistrellus* or *P. pygmeus*, and were therefore excluded from validation. The detector performed well on the validation set, which balanced recall (how well the model finds calls) and precision (how many detections are correct) (see Figure 2) and showed no strong bias towards missing certain species. The classification model also performed well, with most errors being due to confusion of *P. nathusii* and *P. pipistrellus* or assigning false positives to ENV or Myotis, likely due to low frequency and click-like noise both confusing the call detector and classifier.

**Figure 2.**
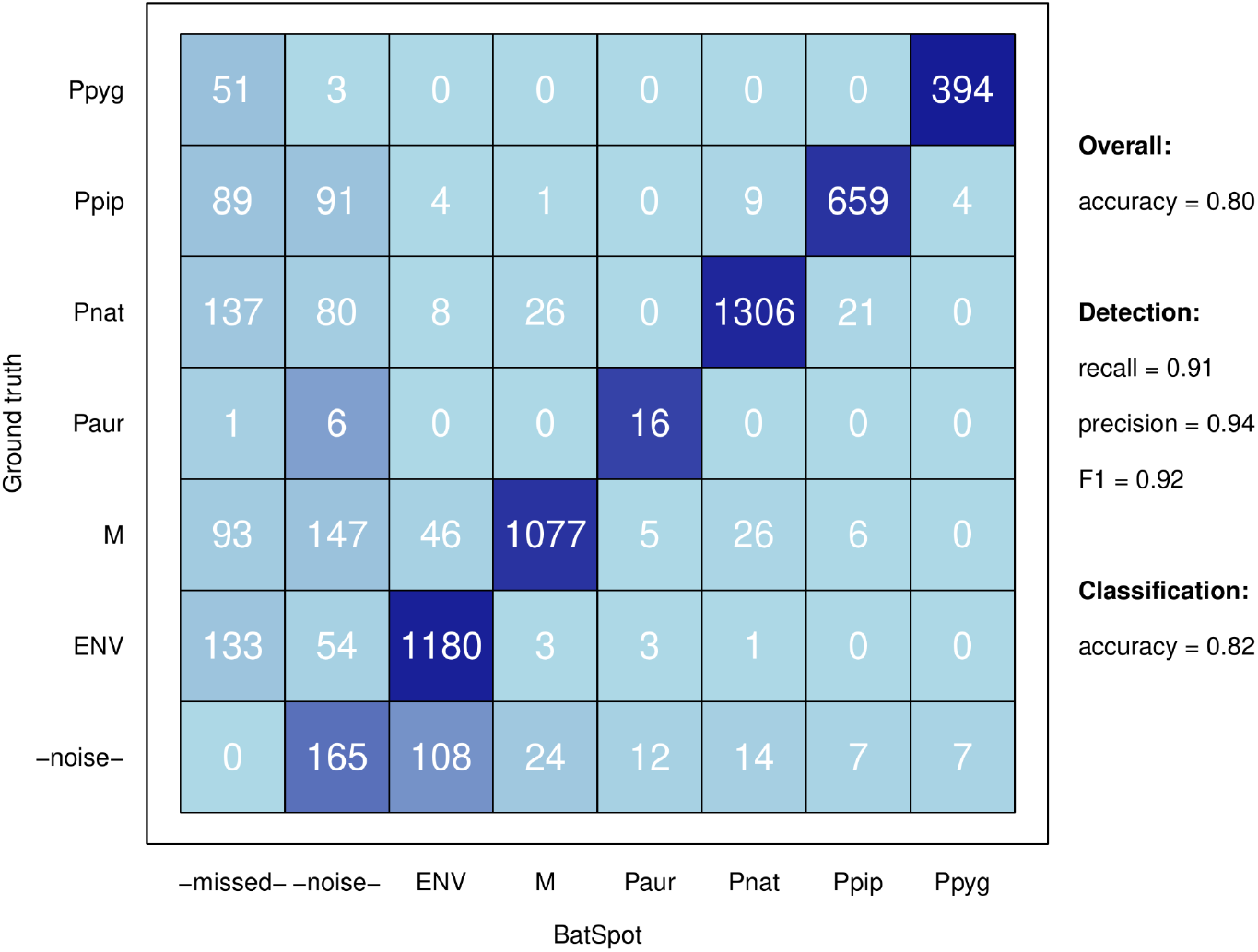
Overall confusion matrix for the call detector and classifier. Rows are the manual annotations, columns are the results from the models for each call. Ppyg = Pipistrellus pygmeus, Ppip = P. pipistrellus, Pnat = P. nathusii, Paur = Plecotus auritus, M = Myotis sp., ENV = Eptesicus serotinus/Nyctalus noctula/Vespertillio murinus, -noise- = calls annotated as noise or classified as buzz/approach/social/Bbar, -missed- = calls not detected by the call detector.

Overall BatSpot outperformed all other tested software (see Figure 3 and 4). BatDetect2 and the BTO bioacoustics pipeline came close in balancing the recall and precision as well as accurately classifying the files with detections (see Table 1).

**Figure 3.**
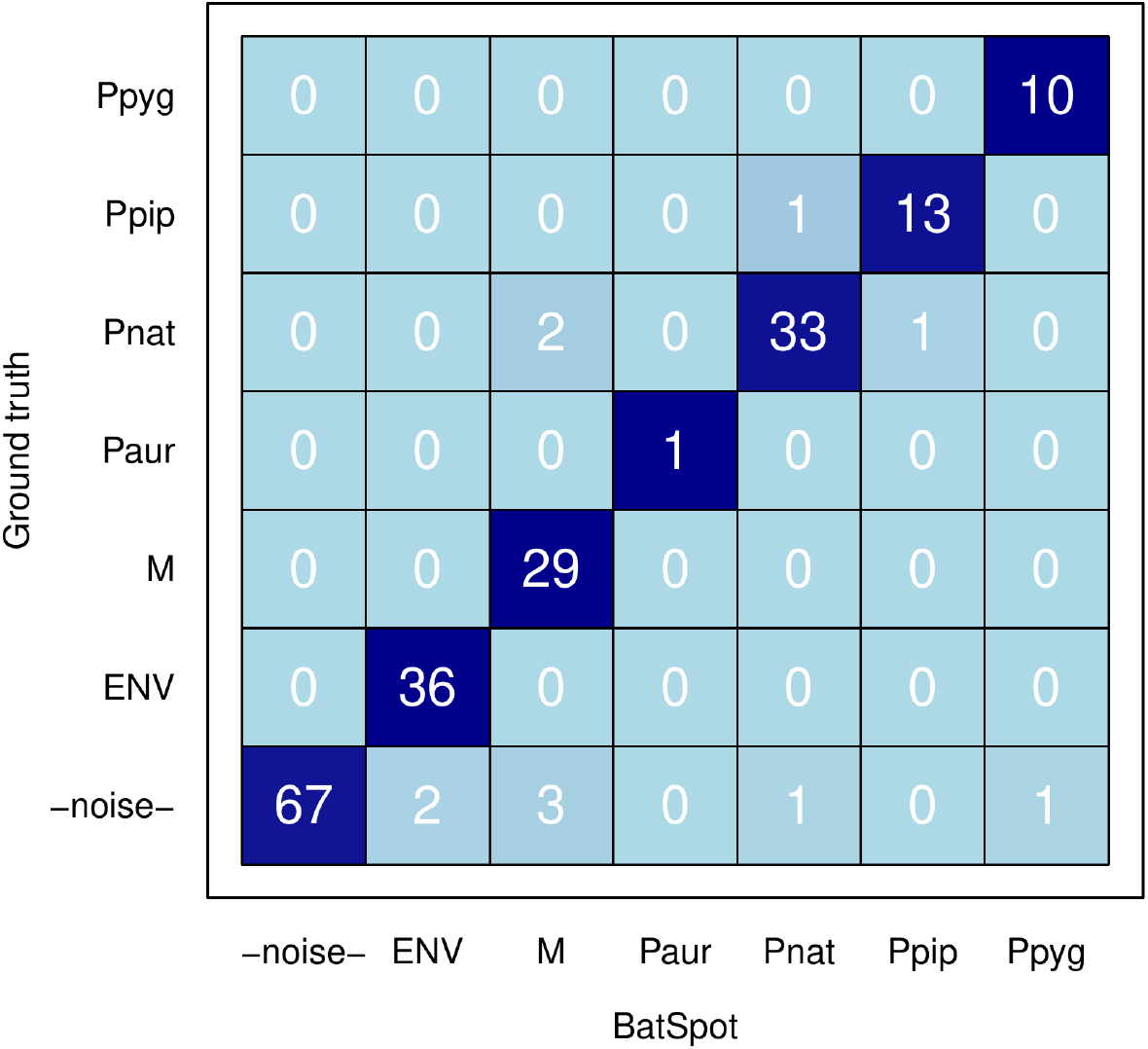
Overall confusion matrix at file level for the call detector and classifier. Rows are the manual annotations, columns are the results from the models for each call. Ppyg = Pipistrellus pygmeus, Ppip = P. pipistrellus, Pnat = P. nathusii, Paur = Plecotus auritus, M = Myotis sp., ENV = Eptesicus serotinus/Nyctalus noctula/Vespertillio murinus, -noise- = calls annotated as noise or classified as buzz/approach/social/Bbar, -missed- = calls not detected by the call detector.

**Figure 4.**
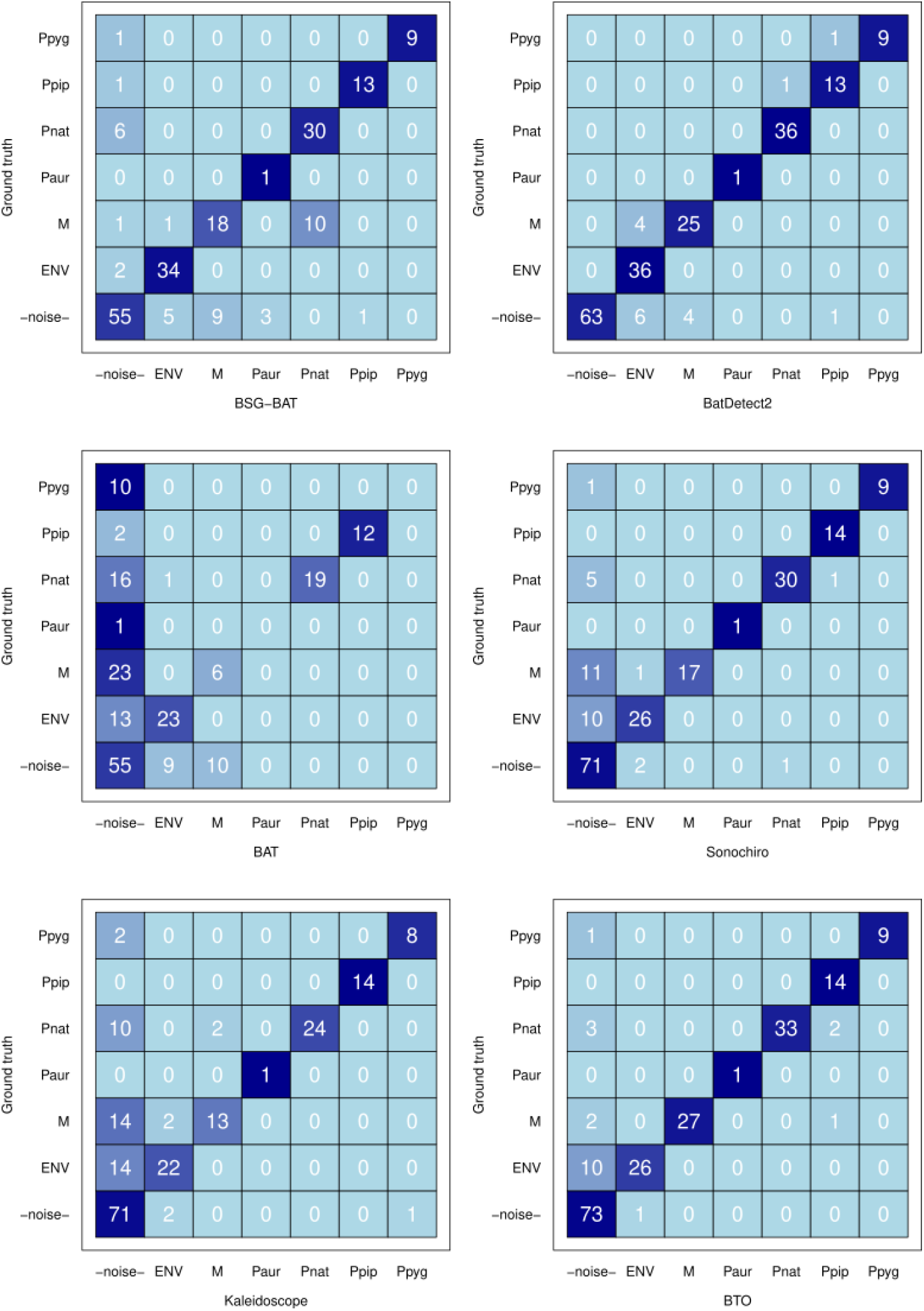
Confusion matrices for BSG-BAT, BatDetect2, BAT, Sonochiro, Kaleidoscope and BTO bioacoustics pipeline. Rows are the manual annotations, columns are the results from the different software at file level. Ppyg = Pipistrellus pygmeus, Ppip = P. pipistrellus, Pnat = P. nathusii, Paur = Plecotus auritus, M = Myotis sp., ENV = Eptesicus serotinus/Nyctalus noctula/Vespertillio murinus, -noise- = files annotated as containing noise or any other bat species.

**Table 1.** Performance of all tested software on the Danish validation set. Recall = tp/(tp+fn), precision = tp/(tp+fp), F1 = 2 * (recall*precision)/(recall+precision) and accuracy = (tp+tn)/(tp+tn+fp+fn) for all files with detections; where tp = number of true positives, tn = number of true negatives, fp = number of false positives and fn = number of false negatives. The highest value(s) in each column is/are highlighted in bold.

|  | Detection |  |  | Classification |
| --- | --- | --- | --- | --- |
| Software | Recall | Precision | F1 | Accuracy |
| BatSpot | **1.00** | 0.95 | **0.97** | **0.95** |
| BSG-BAT | 0.91 | 0.87 | 0.89 | 0.80 |
| BatDetect2 | **1.00** | 0.92 | 0.96 | 0.92 |
| BAT | 0.48 | 0.76 | 0.59 | 0.58 |
| Sonochiro | 0.79 | 0.97 | 0.88 | 0.84 |
| Kaleidoscope | 0.68 | 0.97 | 0.80 | 0.77 |
| BTO | 0.87 | **0.99** | 0.92 | 0.92 |

### Buzz detector

After 100 epochs (the maximum) the model had a validation accuracy of 0.97 and a test accuracy of 0.96. For the validation recordings performance was generally very high (F1: 0.95, recall: 0.96, precision: 0.93). Performance was highest for the Panamanian recordings (F1: 0.97, recall: 0.95, precision: 0.99), intermediate for the Danish recordings (F1: 0.88, recall: 0.85, precision: 0.92) and lowest for the German recordings (F1: 0.84, recall: 0.94, precision: 0.77). Despite the increased threshold for accepting detections (0.9 instead of 0.5), the model detected a lot of false positives for the German recordings.

*Buzzfindr* had much lower performance on the same validation data (F1: 0.37, recall: 0.28, precision: 0.54).

### Social call detector

After 150 epochs (the maximum) the model had a validation accuracy of 0.99 and a test accuracy of 0.98. For the validation recordings performance was lower but still good given the limited dataset (F1: 0.91, recall: 0.93, precision: 0.98).

### Retraining

The call detector trained on only Danish data did not perform very well on the German validation set (F1: 0.48, recall: 0.88, precision: 0.33). Retraining clearly improved the precision (F1: 0.79, recall: 0.79, precision: 0.78) and led to slightly better performance than training from scratch (F1: 0.73, recall: 0.83, precision: 0.64).

## Discussion

Our aim was to develop a tool that can both detect the three most common vocalisation types of bats (search phase calls, feeding buzzes and social calls) and classify search phase calls, the vocalisation type most commonly used for species identification and monitoring studies (Schnitzler & Kalko, 2001; Russo et al., 2018). We present pre-trained models with reasonable out-of-the-box performance and the option to retrain the models with minimal time investment. Finally, we show that the tool can also be used in the Global South. The three detection models presented achieved good performance for the situations that they had been trained for. The buzz detector in particular showed that a single model could detect buzzes in mixed habitats recordings in Denmark, Germany and Panama, with very different noise backgrounds and species compositions.

When applying the search phase call detector model trained exclusively on Danish data to German recordings, we observed a clear drop in performance. This is to be expected, since the noise sources can be very different, even within the same country. This is also clearly what drove the drop in performance, with precision (the proportion of detections that contained search phase calls) dropping much more than recall. Of course, if the detector is used in a country with not only new species (as in Germany), but also new genera or even families that produce search phase calls that do not resemble any of the training examples, a drop in recall is also to be expected. In both cases, retraining will be needed until a dataset with enough examples across the globe becomes available and a global bat detector can be trained. Our hope is that the data presented with this paper can contribute to such a global dataset, and that until a global model is a reality, our Danish model can serve as a basis model to retrain detectors for new locations.

Classification is a more challenging task, and one where even experts with decades of experience cannot always assign call sequences of good quality to a species (Jennings et al. 2008; Rydell et al. 2017). This is simply because some species produce very similar calls when flying in similar environments, and because a given species is often able to modify calls when flying in a different environment. We therefore opted to follow the standard procedure of grouping calls from *Myotis* species into a single category and also grouping calls from species producing lower frequency calls into an ENV species-complex. The classification model was trained with each species separately (with the exception of *M. brandtii* and *M. mystacinus* which could not be separated when generating our training data) to allow the model to learn as many distinguishing features as possible. The grouping was then done after the prediction step, since performance within groups was low as expected. Although an overall accuracy of 0.80 seems low, it should be kept in mind that this includes detection and classification of even the faintest of calls. For many projects (e.g. environmental impact assessments), the aim is to detect and classify species at the file level (as presence/absence in that file), and this is where performance is much higher (F1 = 0.97).

There are also projects (e.g. those aiming to describe the characteristics of calls under varying conditions), in which exact detection of start and end times for individual calls is important. Additionally, when detecting buzzes or social calls, one might want a very low false negative rate, to ensure most target events are detected. This can be accomplished by lowering the detection threshold and subsequently validating all detections manually to get rid of the false positives. Alternatively, one might want a low false positive rate, something particularly relevant when many calls are produced in a sequence and post-hoc processing can be done on the detections. One way to do this is to set up a decision tree, in which, for example, a file is only considered to contain calls from a species if the classifier assigns at least five calls to that species within a sliding window of 3 seconds. This eliminates any false positives that occur spaced out in time.

BatSpot offers a GUI and README for how to retrain, to facilitate obtaining a model with good performance in new recording situations, which is an important step forward and allows researchers to optimise the models without coding experience. An example of this is the offshore environment, where boat, buoy and water noise create challenges to both the detection and classification step by introducing new noise types. With the increase of wind farm development offshore, and the need for both pre- and post-construction monitoring, models performing well in the offshore environment are particularly important. This is one of the reasons that the search phase call detector in BatSpot was trained with a large portion of examples from buoys and wind turbines in the Danish North Sea. That said, BatSpot is far from the first neural network to detect and classify search phase calls, although to the best of our knowledge, it is the first to additionally detect buzzes and social calls. BatDetect2 is a very promising alternative for search phase calls, and achieves comparable performance out-of-the-box. One potential reason for BatDetect2’s high performance is the inclusion of a self-attention layer, which enables the model to integrate information of multiple calls in a sequence to predict the species producing individual calls. With the integration into a simple python package, BatDetect2 requires minimal coding experience, and might therefore be the best option if there is no time for targeted model retraining.

For buzzes and social calls, BatSpot presents a much bigger step forward. The detection of buzzes is important when looking at foraging behaviour, but has previously received almost no attention in the literature. The R package *buzzfindr* (Jameson, 2024) is an important step forward, but struggles to detect faint buzzes and buzzes in noisy environments, something important in settings with multiple individuals flying in an environment with much insect noise as well (as is the case for many of the German and Panamanian recordings). The fact that a single model was able to perform well in three countries across two continents is a major step forward. Social calls have received even less attention and are also particularly difficult to detect automatically, because this category basically includes all vocalisations not used to detect the surroundings. With our current model in BatSpot, it should be possible to reliably detect longer social calls, such as the trills produced by species we have evaluated in the *Pipistrellus* genus. The data presented with the paper will also serve as an important contribution to a future comprehensive dataset with recordings from across Europe or wider. We envision that the social call detector, once further developed, will be able to support passive acoustic monitoring purposes by enabling automated analysis and localisation of important roosting and mating areas.

For all three models, there are several steps to keep in mind when applying them to large datasets. First, a validation set is necessary to quantify the recall and precision for each location and season included in the dataset to be analysed. This does not need to be a specific percentage of all annotated data, but should rather ensure that all variation in the full dataset is covered and performance on it can be quantified. Validating performance is not specific to BatSpot, as even established commercial software might show a strong bias when applied in, for example, the offshore environment (Smeele et al. 2026). If retraining is needed, this can be done by following the steps in the README (see below). This should drastically reduce training time compared to training with all the published data plus the data collected in the specific project (Verwimp et al., 2025). And finally, once the model performs satisfactorily, all raw data can be processed using the scripts supplied to index and process large datasets on a high performance computing cluster, or a local computer with GPU.

## Conclusions

In this article we presented BatSpot, a set of convolutional neural networks to detect search phase calls, buzzes and social calls of bats as well as classify the search phase calls.

Interaction with the models is facilitated through a GUI, but the modified ANIMAL-SPOT source code is still in place, allowing advanced users to run models from the terminal as well. We show that the search phase call detector can be retrained with minimal effort and that the buzz detector works well across three countries and two continents. Overall, BatSpot will enable researchers with no coding experience to tap into the flexibility and power of convolutional neural networks, while at the same time contributing to a growing open source set of annotated recordings, which can be used to train globally applicable models.

